# Exposure to complex mixtures of urban sediments containing Tyre and Road Wear Particles (TRWPs) increases the germ-line mutation rate in *Chironomus riparius*

**DOI:** 10.1101/2024.10.17.618802

**Authors:** Lorenzo Rigano, Markus Schmitz, Volker Linnemann, Martin Krauss, Henner Hollert, Markus Pfenninger

## Abstract

Tyre and road wear particles (TRWPs) are a significant yet often underestimated source of environmental pollution, contributing to the accumulation of microplastics and a complex mixture of contaminants in both terrestrial and aquatic ecosystems. Despite their prevalence, the long-term evolutionary effects of TRWPs, beyond their immediate toxicity, remain largely unknown. In this study, we assessed mutagenicity in the non-biting midge *Chironomus riparius*, upon exposure to urban sediment collected from a runoff sedimentation basin. To assess the extent of mutagenic effects over multiple generations, we combined the urban sediment exposure model with short-term mutation accumulation lines (MALs) and subsequent whole genome sequencing (WGS). Our results reveal that the exposure to urban sediment significantly increases mutation rates compared to control groups by 50%, independent of concentration (0.5% and 10%). To infer potential causal processes, we conducted a comparative analysis with known mutational spectra from experiments with other studies. This comparison showed that the mutation profiles induced by urban sediment clearly clustered with those caused by Benzo[a]Pyrene (BaP), a known polycyclic aromatic hydrocarbon (PAH). A comprehensive chemical characterization of the sediment confirmed a considerable impact of road runoff and traffic-related contamination, including PAHs of primarily petrogenic origin. This suggests that PAH-like compounds present in urban sediments may play a significant role in the observed mutagenic effects. Our study shows that urban sediments influence mutation rates and alter mutational spectra in exposed organisms, potentially compromising genomic stability and shaping evolutionary trajectories. Additionally, we show that comparatively analysing mutational spectra may provide valuable insights into mutational processes. These genetic changes may have profound long-term effects on population dynamics and ecosystem health, underscoring the importance of understanding the evolutionary consequences of environmental pollution.

## 1. Introduction

Globalisation is largely driven by an ever-expanding transport network of people and goods, transforming global trade and mobility. However, this progress comes with multiple costs (Sovacool et al., 2021). One of the less recognised consequences of road transport are Tyre and Road Wear Particles (TRWPs). As vehicles travel on asphalt surfaces, the inevitable wear generates constantly particles. These tiny fragments, ranging from nanometers to micrometers in size, consist of a mixture of rubber, asphalt components, and other various environmental contaminants. The widespread presence of TRWPs in water bodies, soils and airborne particles highlights their environmental mobility (Baensch-Baltruschat et al., 2021; Sieber et al., 2020; Unice et al., 2019), raising concerns about their ecological impact. In Germany alone, approximately 20,000 tonnes of TRWPs enter water bodies annually, representing one of the largest sources of microplastics released into the environment (Baensch-Baltruschat et al., 2021). Once microplastic particles enter aquatic ecosystems through rainwater runoff or wind, they can harm organisms in these environments (Khosrovyan et al., 2022).

One of the major entry points for TRWPs into aquatic ecosystems is urban runoff, which is managed by sedimentation basins strategically placed near busy roads. Their primary role is to separate water and particles from road runoff through sedimentation, allowing the purified water to enter nearby water sources while retaining the sediments within the basins (Grung et al., 2016). However, road runoff often enters water bodies or surrounding areas untreated (Grung et al., 2016), making sediment accumulated in urban runoff sedimentation basins an ideal subject to assess the ecotoxicological effects of TRWPs and urban particulate matter.

The introduction of anthropogenic substances into the environment can adversely affect organisms by lowering their fitness. In this case, the alteration of the environment caused by these stressors modifies the selective pressures acting on them (Charlesworth, 2009, 1971). The key factor for the evolutionary response to anthropogenic pollution is therefore their effect on population fitness, which on the one hand integrates all processes within individuals and on the other hand is decisive for the organism’s role in the ecological community and thus for the ecosystem (Sæther and Engen, 2015; Shaw and Geyer, 2010). Adaptation to adverse changes in the environment by environmental pollutants is widespread and can occur rapidly within a few generations (Barghi et al., 2020; Bulut et al., 2024). Changes in population fitness may be associated with changes in demography (Shaw and Geyer, 2010; Tuljapurkar, 1982), which in turn may affect other species in the community, thus triggering eco-evolutionary dynamics (Carroll et al., 2007; Fussmann et al., 2007).

Another impact of anthropogenic stressors on evolutionary fate is the induction of mutations that, although essential for evolution, in most cases have a negative impact on fitness (Eyre-Walker and Keightley, 2007). Therefore, anthropogenic substances should be considered not only as drivers of immediate lethal and sublethal toxic effects but also as potential agents of natural selection that may trigger profoundly change the evolutionary trajectory (Ankley et al., 2010; Brady et al., 2017). However, current ecotoxicological research tends to focus more on the immediate ecological impacts of tested contaminants, rather than their perhaps even more and certainly irreversible evolutionary consequences. Thus, to fully understand the impact of anthropogenic stress on biodiversity, it is essential to study its evolutionary effects on organisms.

*In vitro* studies have demonstrated that TRWPs can induce cytotoxicity through various biological mechanisms, with inflammation and oxidative stress being the most evident (Fussell et al., 2022). Furthermore, genotoxicity has been reported as another mechanism of toxicity, as DNA damage has been observed after exposure of human lung cells to TRWPs (Bouredji et al., 2023). Although genotoxicity, which refers to damage to DNA in somatic cells, may occur, it does not necessarily result in mutagenicity, meaning it does not always induce mutations in germ line cells that can be inherited and thus have implications for evolutionary processes (Doria and Pfenninger, 2021).

In our study, we investigated the microevolutionary responses of *Chironomus riparius* to urban sediments from a runoff sedimentation basin by assessing mutagenicity across multiple generations. Since oxidative stress is a known trigger of mutations (Aitken and Krausz, 2001), we measured ROS levels to evaluate the extent of oxidative stress. We used a recently introduced mutation rate test (Oppold and Pfenninger, 2017) as an effective ecotoxicological mutagenicity test tool for metazoan organisms (Doria and Pfenninger, 2021). Previous studies, in which *C. riparius* was exposed to Cadmium (Cd) and Benzo[a]pyrene (BaP), substances known for their potential mutagenicity, have shown the robustness of our approach in assessing germline mutagenicity (Bulut et al., 2024; Doria and Pfenninger, 2021). The test consists of a combination of short-term mutation accumulation (MA) lines, whole genome sequencing (WGS), and dedicated data analysis. After exposure to potentially mutagenic substances, whole genome sequencing (WGS) of individuals from these lines can reveal *de novo* mutations compared to the initial parents. By analysing the number and spectrum of these mutations, the mutagenic potential of the tested compound can be investigated. In addition, we introduce a comparative analysis of the mutational spectra induced on *C. riparius* by different environmental stressors.

The non-biting midge *C. riparius* is an established model organism in evolutionary ecotoxicology (Foucault et al., 2019) of which we have the fully sequenced genome on chromosome scale (Pettrich et al., 2024; Schmidt et al., 2020), enabling detailed analysis of mutations and their potential functional consequences. Additionally, *C. riparius* has a relatively short generation time, allowing efficient propagation of MA lines and rapid detection of mutagenic effects. Furthermore, as a common inhabitant of freshwater ecosystems, *C. riparius* is frequently exposed to contamination from urban runoff in nature and thus a realistic test organism.

## 2. Materials and Methods

### 2.1 Culturing conditions

The culture of *C. riparius* used in the present study came from a local population collected in a small river located in Hasselbach, Hessen, Germany (co-ordinates: 50.167562N, 9.083542E). This culture was maintained and regularly replenished from the field as an in-house laboratory standard. The cultivation conditions of the stock culture conformed to a modified protocol based on the methodology outlined in OECD guideline N219, as previously described (Foucault et al., 2019).

### 2.2 Sediment collection and preparation

In February 2023, the sediment was collected from an urban runoff sedimentation basin located in Aachen, Germany (50°48’03.6”N 6°06’29.7”E) receiving runoff via a separate sewer system from a large area in the district of Aachen Soers, including a sports park and a highly frequented road, Bundesstraße B57 Krefelder Straße. The total drained area covers 68 hectares. The B57 serves about 26.000 vehicles a day (BASt, manual counting; 2021). The sampling took place during cleaning of the basin, and samples were obtained from different spots within the basin using a stainless-steel bucket, thoroughly mixed, and stored in a cold room at 4 °C until further use. For further analyses, aliquots were freeze-dried and sieved (stainless steel mesh, 1 mm mesh size) to remove large particles at the Goethe University, Frankfurt am Main.

### 2.3 Mutation accumulation lines (MALs) experimental design

All MALs originated from a single egg rope (F0) raised under optimal conditions to avoid selection. Subsequently to the successful reproduction of its offspring (F1), forty-five egg-ropes were collected and used to establish 65 MA lines. The hatched clutches were placed individually in glass bowls (∅20 × 10 cm) containing three increasing concentrations of urban sediment, as a percentage of total weight:

The quartz playground sand* used had a granularity of 0-2 mm, was pH neutral, and was washed before use. In the glass bowls containing the urban sediment, the sand and the urban sediment were mixed, so that the resulting sediment column was homogenised. To all glass bowls, 1.250 L of medium was added. Only a single egg-clutch originating from each MA line was chosen to start the following next generation. The test vessels were maintained in a constantly ventilated room at a temperature of 20 ± 1°C, with a relative humidity of 60% and a 16:8-hour light/dark cycle. To compensate for water evaporation in the test vessels, demineralized water was constantly supplemented. Conductivity was maintained between 550 and 650 mS/cm and pH around 8. The replicates were then raised and fed daily with finely ground fish food (Tetramin® Flakes) whose amount was calculated for the respective developmental stage as described elsewhere (Foucault et al., 2019).

Because of the swarm fertilization of *C. riparius* and the thereby following impossibility of determining the parents of specific egg clutches, adult offspring of the first generation were collected and used as a pooled reference to infer the joint genotype of the parents. At the end of the fifth generation, one randomly chosen female was collected for DNA extraction.

### 2.4 Whole Genome Sequencing and Bioinformatic analysis

DNA extraction was performed with the Blood and Tissue QUIAGEN Kit following the manufacturer’s instructions. The whole genome sequencing of both the reference pool and individuals was established using the study (Doria and Pfenninger, 2021). Clean reads of individual females of each mutation accumulation line and ancestors were analyzed using the best practices of the GATK pipeline (McKenna et al., 2010). First, the reads for POOL were paired with Pear (Zhang et al., 2014). The reference genome v.4 (unpublished data) was used for mapping with bwa-mem. Picard v.1.123 (https://broadinstitute.github.io/picard/) was used for marking and removing duplicates, and low-quality reads were removed using samtools with default parameters. The target lists for realignments, vcf file creation, variant filtration, and base recalibration were created using GATK. The bam files were merged with samtools merge (Danecek et al., 2021), and accuMUlate (Winter et al., 2018) was used with the same reference genome and merged bam files. The output was filtered with a custom bash script using the following parameters: probability of a mutation (≥ 0.90), probability of one mutation (≥ 0.90), probability of correct descendant genotype (≥ 0.90), N mutant in wt (= 0), mapping quality difference (≤ 2.95), and stand bias (≥ 0.05). The filtered mutation positions were then manually validated using IGV. The mutation rate was estimated by dividing the number of mutations per generational passage times the callable sites.

### 2.5 Chemical analyses

The sediment was screened for Polycyclic Aromatic Hydrocarbon content (16 EPA PAHs) using GC-MS (DIN 38407-39:2011-09), Heavy Metals using ICP-MS (DIN EN 11885:2009-09), as well as mineral oil indices (DIN EN ISO 9377-2:2001-07) measured by the Institute of Environmental Engineering at the RWTH Aachen University in freeze-dried particles.

Moreover, organic extracts were prepared using an exhaustive ultrasonic-assisted extraction via an elutropic solvent gradient adapted from previous methods (Hiki and Yamamoto, 2022; Klöckner et al., 2021). A series of two solvents from n-heptane-acetone (1:1 v/v) and methanol was used, covering a broad spectrum of the elutropic series (Snyder, 1974). 400 mg of sediment (sieved to 1000 µm) was suspended in 10 mL of solvent, sonicated for 15 min. in an ice-cooled water bath to prevent overheating, and then centrifuged at 1900 x G for 2 minutes. After centrifugation, the supernatant was carefully aspirated using a glass Pasteur pipette and transferred to a Büchi round flask. The extraction was repeated once more with n-heptane-acetone mixture and two more times with methanol. After these four extraction steps, the combined extracts were evaporated to almost dryness, redissolved in 1 mL methanol, and filtered using a 0.2 µm PTFE-membrane filter attached to a luer-lock syringe to remove finest aspirated particles. For target-screening on tyre rubber additives and wastewater pollutants the extracts were diluted 5-fold before being analysed using liquid chromatography coupled to high-resolution mass spectrometry (LC-HRMS) at the Department Exposure Science at the Helmholtz Centre for Environmental Research.

For LC-HRMS analysis a Thermo Ultimate 3000 LC system coupled to a quadrupole-Orbitrap instrument (Thermo QExactive Plus) with electrospray ionisation in positive and negative ion mode was used. For LC separation a Kinetex Biphenyl column (100 × 2.1 mm, 2.6 µm particle size) equipped with in-line filter and pre-column of the same type (5x 2.1 mm) was employed. The gradient separation uses 0.1% formic acid (eluent A)/methanol containing 0.1% formic acid (eluent B)/acetonitrile (eluent C) for positive mode and 1 mm ammonium fluoride (A)/methanol containing 1 mM ammonium fluoride (B)/acetonitrile (C) for negative mode. The gradient program started for both modes at 97% A/3% B/0% C, held for 2 minutes, increasing to 3% A/97% B/0% C in 14 minutes, and changing to 3% A/0% B/97% C in 4 minutes, held for 4 minutes. The eluent was afterwards re-equilibrated to initial conditions in 4.8 minutes. The HRMS analysis combined a full scan acquisition (m/z 80-1200, nominal resolving power of 70,000) with six data independent acquisition scans (m/z 80-182, 178-282, 278-382, 378-482, 478-682, 682-1200) at a nominal resolving power of 35,000.

For quantification, Thermo raw files were converted to mzML files using ProteoWizard (Chambers et al., 2012), followed by peak detection and annotation in MZmine 2.38 (Pluskal et al., 2010) and further processing by the MZQuant R package (Finckh et al., 2022). We used an interna l standard calibration with target analyte concentrations ranging from 0.2 to 1000 ng/mL and an internal standard concentration of 50 ng/mL.

### 2.6 ROS Detection and Image Analysis

First-stage chironomid larvae were subjected to the three increasing concentrations of urban sediment (0, 0.5% and 10%). Subsequently, 20 L3 larvae were collected and placed in 24-well plates. Plates were filled with 2.5 ml medium as described before (Foucault et al., 2019). CellROX Orange (Thermo Fisher cat. no. C10443) reagents were used to identify ROS products. CellROX is an oxidative stress reagent that is cell-permeable and suitable for live cell ROS measurements. Within the reduced state, they are non-fluorescent but after oxidation by ROS, they exhibit fluorogenic signals at 545/565 nm for CellROX Orange. The reagent is localised within the cytoplasm and can detect 5 different ROS types (hydrogen peroxide, hydroxyl radical, nitric oxide, peroxynitrite anion, and superoxide anion). After placing larvae on the well plates, 0.75 µl of CellROX Orange was used per larva.

Well-plates were placed in a climate chamber with a 16:8 light/dark cycle with 550 lux light intensity without aeration under 20°C. After 24 hours of treatment, well plates were placed in a styrofoam box to avoid the temperature change effect. The ROS was measured in a live larva with ZEISS Axio Imager 2 under 10x magnification. The images were taken with AxioVision Rel. v.4.8. For fluorescence images, an HXP 120 C fluorescence lamp was used with maximum light intensity (Item Number: 423013-9010-000). Fluorescence images were obtained from the larva under filter set “43 HE” (BP 550/25 HE, FT 570 HE, BP 605/70 HE, Item Number 489043-9901-000) with 1 second exposure. This specific filter excites blue light around 550 nm, transmitting emitted red fluorescence above 570 nm filtering out the remaining blue excitation light and allowing only red fluorescence around 605 nm.

The fluorescence field images were analysed by ImageJ Fiji (v. 2.15.0). Images were uploaded to ImageJ as an image sequence and converted to 8-bit grayscale from RGB Color images to avoid colour difference and only calculate light intensity. The same threshold was applied to all images (Threshold: 23). After setting the threshold measure function was used. The mean values of each image were taken as fluorescence intensity. The fluorescence intensity we measured is not the actual ROS amount within the organism, but the current amount of reagent entered in the organism and oxidised by binding to ROS, which is the remaining amount after the organism’s antioxidant system scavenges ROS.

### 2.7 Heatmap and dendrogram of different environmental conditions

To compare mutational spectra of different environmental conditions from previous experiments, we constructed a heatmap and a dendrogram based on the similarity of mutation rates for the different mutational classes of *C. riparius*. Data were collected from 14 experimental conditions, covering six classes of nucleotide substitutions (G<>T, A<>T, C<>G, A<>C, C<>T, G<>A). The heatmap, generated with ggplot2, displayed experimental conditions on the y-axis and nucleotide substitutions on the x-axis. A grayscale gradient, ranging from light gray (low values) to black (high values), was used to represent transition and transversion rates, emphasizing differences in mutation rates between conditions. Clustering patterns were emphasized through faceting to clearly show trends. The dendrogram was created using hierarchical clustering with Ward’s method on a distance matrix derived from scaled mutation data, illustrating the similarity between experimental conditions based on their mutational spectra. All data analysis, clustering, and visualization were performed using R (v. 4.2.1).

### 2.8 Statistical analysis

Data from ROS measurements were analysed using the R package brms to perform a Bayesian-ANOVA, to test the differences between treatments. The analysis included Bayesian linear regression to model the treatment effects, and posterior distributions along with credible intervals were computed to assess the differences in means and their significance. To compare mutational rates between treatments, a Bayesian Poisson test was applied using the R package BayesianFirstAid (Bååth, 2014).

## 3. Results

### 3.1 Mutation rate and spectrum

Mean sequence coverage ranged from 31.8x to 37.9x for the individual MALs lines and 55.7x for the reference pool. The number of callable sites ranged from 137,847,045 bp to 161,044,682 bp. Overall, we identified 27 mutations during 60 generational passages in control, 38 mutations during 70 generational passages in 0.5% and 42 mutations during 70 generational passages in 10% (Table 2). For the control, 7 transversion and 20 transition mutations were identified. In 0.5% urban sediment, 17 transversion, and 21 transition mutations were identified. In 10% urban sediment, 13 transversion, and 29 transition mutations were identified (Table 3).

**Table 1:** The three increasing concentrations of urban sediment tested.

| 0 (control) | 0,5% | 10% |
| --- | --- | --- |
| 500 g of quartz playground sand* | 497,5 g* + 2.5 g of urban sediment** | 450 g* + 50 g** |

**Table 2:** Mean coverage, number of callable sites, number of single nucleotide mutations (SNM), and mutation rate (μ) per treatment of *C. riparius*.

| Treatment | Mean coverage | N. of callable sites | N. of SNM | Mean $\mu$ Rate |
| --- | --- | --- | --- | --- |
| Control | 33.1 | 1.6E+08 | 27 | 2.79E-09 |
| 0.5% urban sediment | 31.8 | 1.4E+08 | 38 | 4.11E-09 |
| 10% urban sediment | 37.9 | 1.5E+08 | 42 | 4.23E-09 |

**Table 3:**
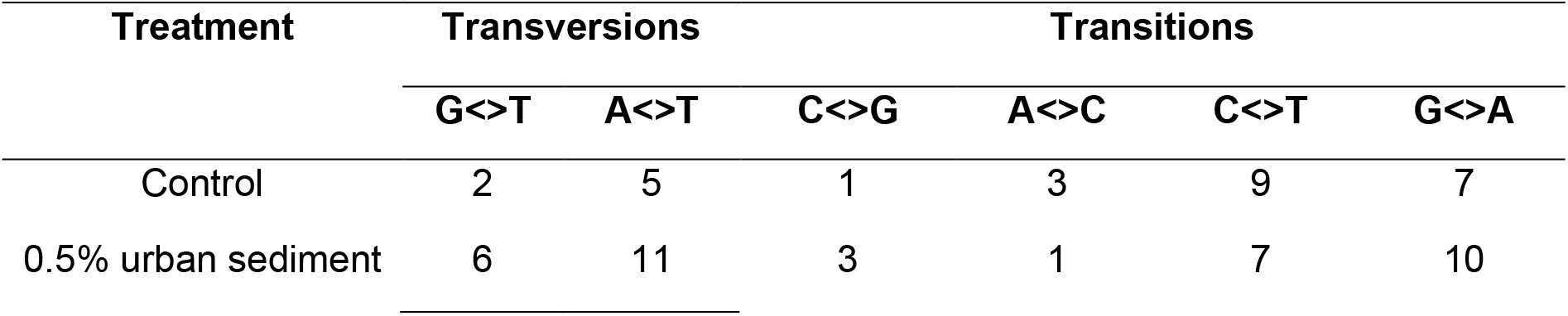
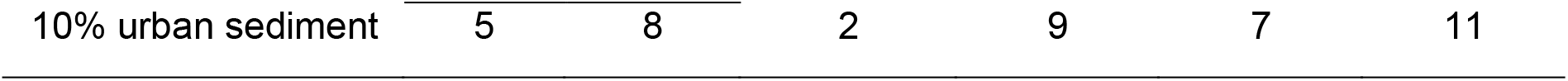
Mutation spectrum of *C. riparius* after 5 generations of urban sediments exposure.

The mutation rate estimate for the control was μ = 2.79 × 10^−9^ (95% HDI 1.8 × 10^−9^ and 4.0 × 10^−9^), for 0.5% of urban sediment was μ = 4.11 × 10^−9^ (95% HDI 2.9 × 10^−9^ and 5.5 × 10^−9^) and for 10% of urban sediment was μ = 4.23 × 10^−9^ (95% HDI 3.1 × 10^−9^ and 5.6 × 10^−9^). The 0.5%/control rate ratio was 1.5, with 93.9% certainty this ratio being larger than 1. The 10%/control rate ratio was 1.5 (95.5% certainty). The 0.5%/10% rate ratio was 0.97 (55%<1<45%) (Figure 1).

**Figure 1:**
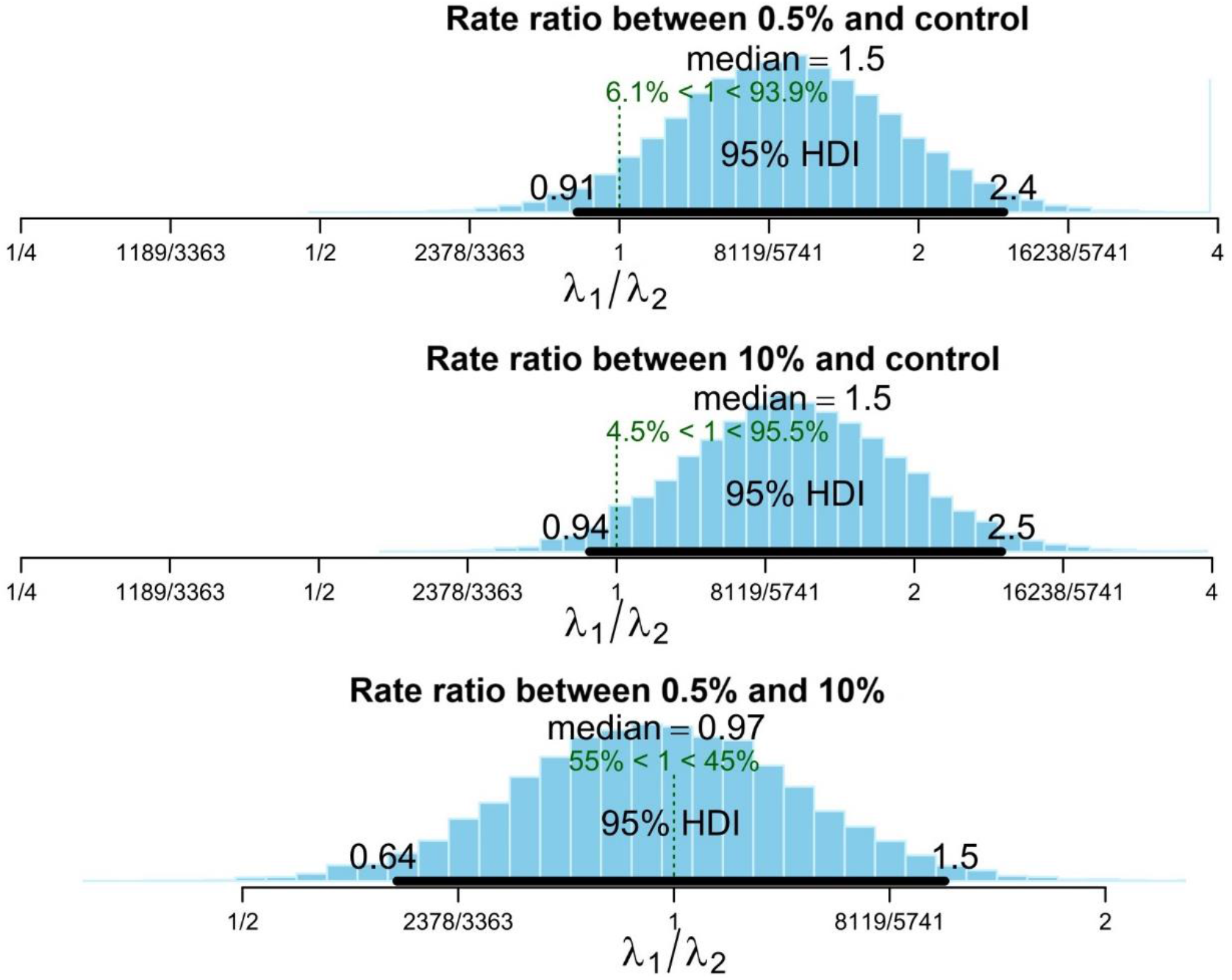
Posterior distributions for rate ratios between treatment groups.

### 3.2 Chemical characterisation of urban sediment

The chemical characterisation of the sediment from the investigated lamellar basin revealed a broad spectrum of anthropogenic and in particular urban and traffic-related contamination (Figure 2). The total organic carbon content of the investigated sediment was 124 g/Kg sediment dry weight of which a maximum measured tyre rubber content (TWPmax) was 5.2 g resulting in a total share of 0.52% tyre rubber. Based on TWPmax measurements in road runoff from highly frequented roads in Aachen city of 11.5% TWPmax, we can estimate a contribution of 4.5 to 8.5% road runoff to the overall urban runoff share in the sampled lamella basin. The total contamination load, including salts, metals, PAHs, oils, and synthetic chemicals added up to 87.91 g/Kg sediment dry weight. Of these, the majority (52.46%) was shared by road salts with a total of 46.12 g. The second highest contribution was by heavy metals (24.65%) with 21.67 g/Kg dw and Aluminum (Al) (13.07 g/Kg dw; 14.86% of contaminants load). Heavy metal contamination was largely dominated by Iron (Fe) with 19.5 g/Kg dw, followed by Zinc (Zn) (1.74 g/Kg dw; 1.98%) and Copper (Cu) (144 mg/Kg dw; 0.16%).

**Figure 2:**
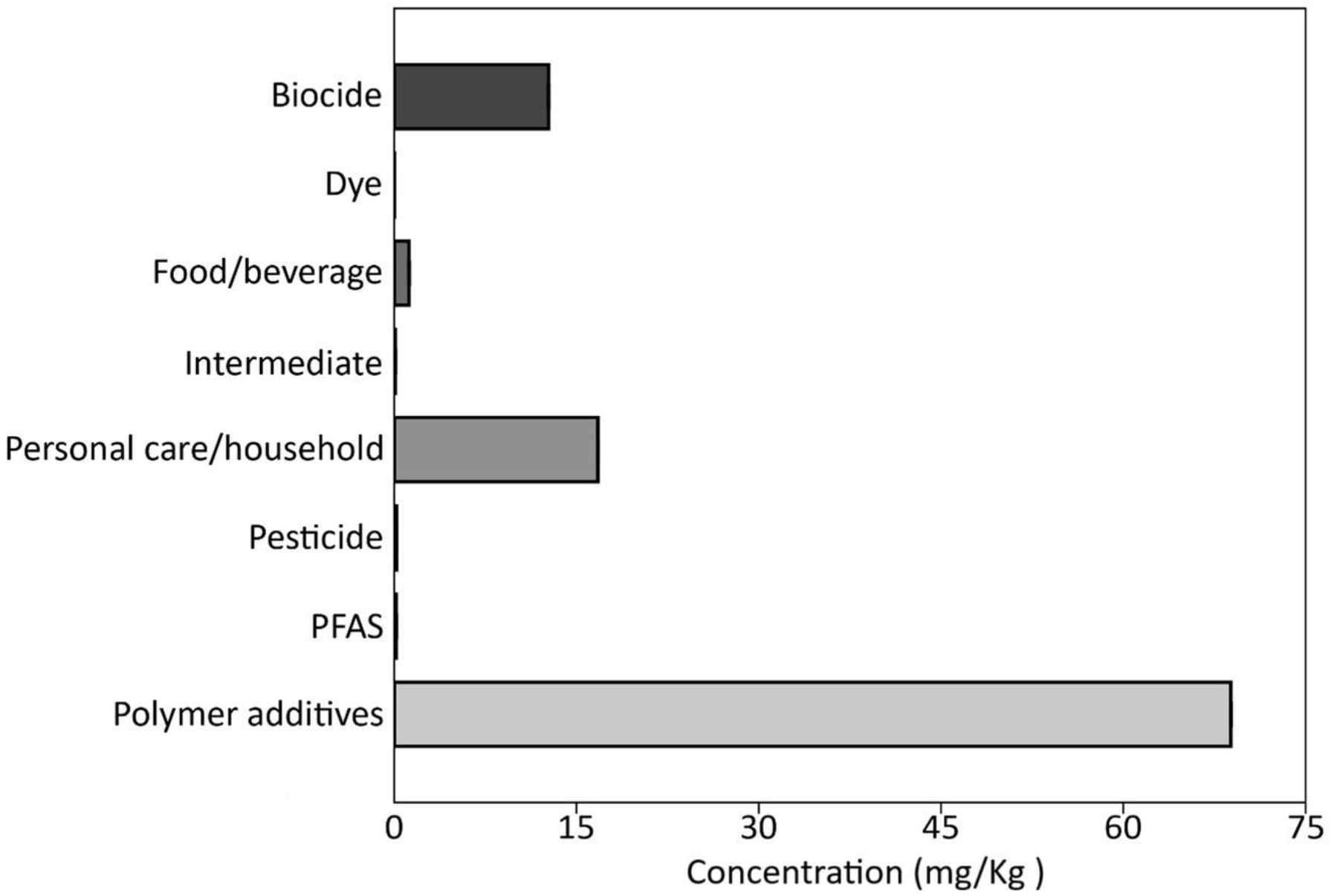
Tyre rubber additives and wastewater chemicals HP-HRMS/MS target screening in lamella basin sludge organic extracts. The sum of concentrations is indicated in mg/Kg sediment equivalent.

The screening of the 16 EPA PAHs returned 4 mg PAHs per Kg sediment of which about 80% were shared by high molecular weight PAHs, such as Benzo(b)fluoranthene (1200 µg/Kg), Benzo(g,h,i)perylene (1090 µg/Kg), and Benz(a)anthracene (633 µg/Kg). Additionally, 983 µg/Kg mineral oils were measured in the sediment.

The measured PAHs content differs from a previously reported sampling of the investigated site. The sediment in this study contained a share of 4.0 mg/Kg dw PAHs, whereas sediment sampled earlier contained 2.9 mg/Kg dw. The Fluoranthene to pyrene ratio is 0 as these were below the LOD (0.55 in Rigano et al. (2024)), the Indeno(1,2,3,c,d)pyrene to Benzo(g,h,i)perylene ratio was 0.14, comparable to our previous study. The considerable lower share of low molecular weight PAHs and resulting altered composition hints to a stronger contribution of PAHs of petrogenic origin (Tobiszewski and Namieśnik, 2012).

The LC-HRMS/MS target screening on wastewater chemicals and tyre rubber additives revealed a strong contribution of known plastic and rubber additives. Of the total 124 detected synthetic chemicals, 49 were labeled as polymer additives, 23 as personal care and household chemicals, 19 as pesticides, 14 as biocides, and 12 as food and beverage related. Additionally, three PFAS compounds, three intermediate compounds, and one industrial dye were detected. Of the top 50 most abundant substances detected (by mass), 31 were identified as tyre rubber additives, composed mainly of phosphate-esters (38,7 mg/Kg dw, n=8) used as flame retardants, phthalates (10,1 mg/Kg dw, n=4) used as plasticisers, benzothiazole derivates (2,1 mg/Kg dw, n=3) used as vulcanization agents, and bisphenols (2,6 mg/Kg dw, n=2), also used in tyre manufacturing. Additionally, the ubiquitous tyre rubber additives N’N-diphenyl-guanidine (3.1 mg/Kg dw), and 6PPD-quinone (112 µg/Kg dw) were detected.

### 3.3 Reactive oxygen species measurements

The fluorescence images were analysed with ImageJ to determine the red colour density and intensity. A difference of means of -7.21 was found between control and 0.5% (95% HDI, between -11.96 and 2.45), with posterior probability of 98.8%, that control was smaller than 0,5%. A difference of means of 2.64 was found between control and 10% (95% HDI, between 1.92 and 7.35), with posterior probability of 100%, that control was larger than 10%. A difference of means of 9.85 was found between 0.5% and 10% (95% HDI, between 5.00 and 14.71), with posterior probability of 100%, that 0.5% was larger than 10% (Figure 3).

**Figure 3:**
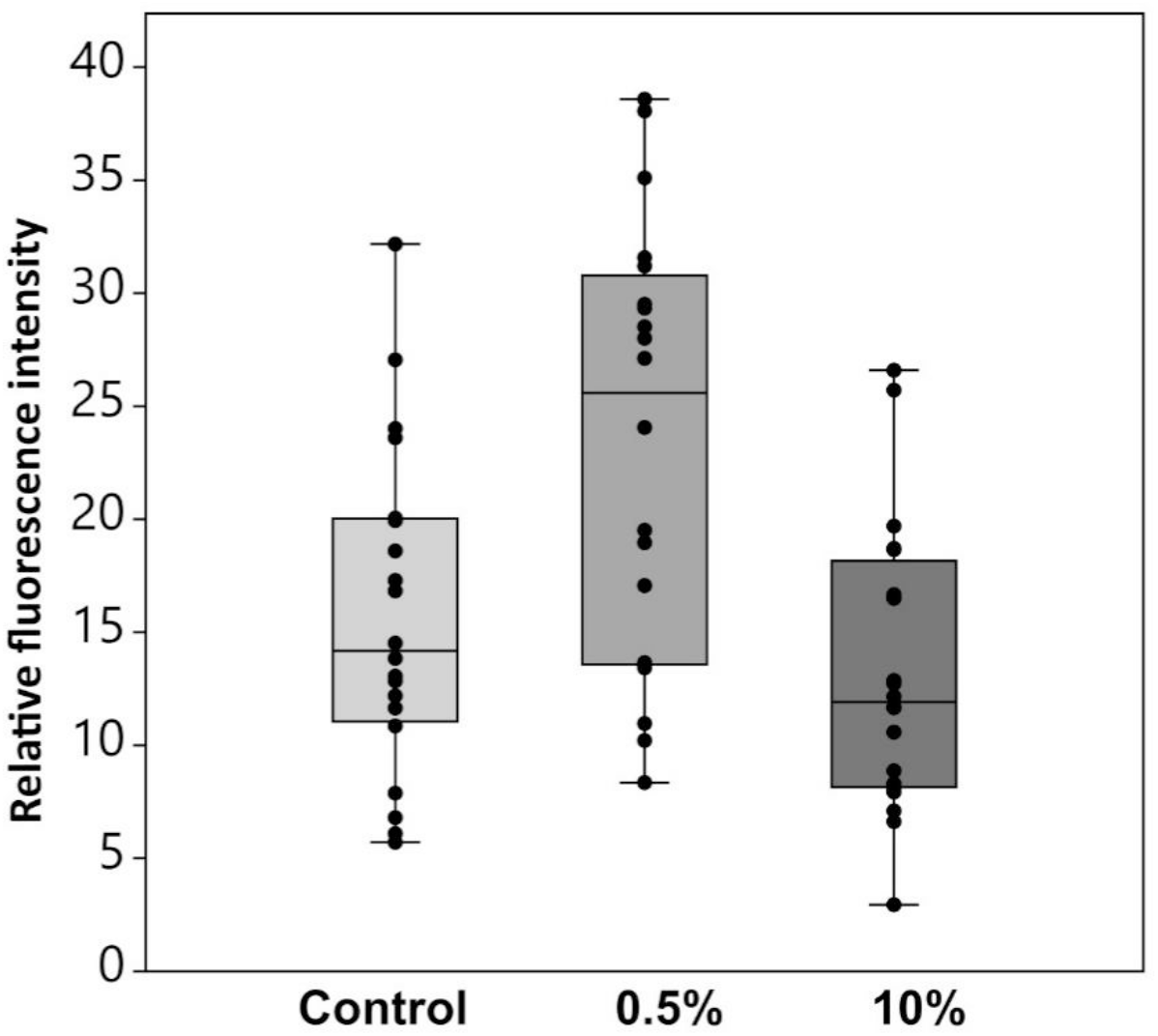
Comparison of relative fluorescence intensity between control, 0.5% and 10%.

### 3.4 Heatmap and dendrogram

The dendrogram results show that the control groups cluster separately from the other treatments. Among the treatments, the results show that, apart from the highest temperature of 26°C, which clusters with Cd, the intermediate temperatures (16°C to 23°C) cluster together. The lowest temperature, 12°C, clusters with the highest concentration of BaP (100 μg/L) and the highest concentration of urban sediment (10%). In contrast, the two lowest concentrations of urban sediment (0.5%) and BaP (10 μg/L) cluster exclusively together (Figure 4).

**Figure 4:**
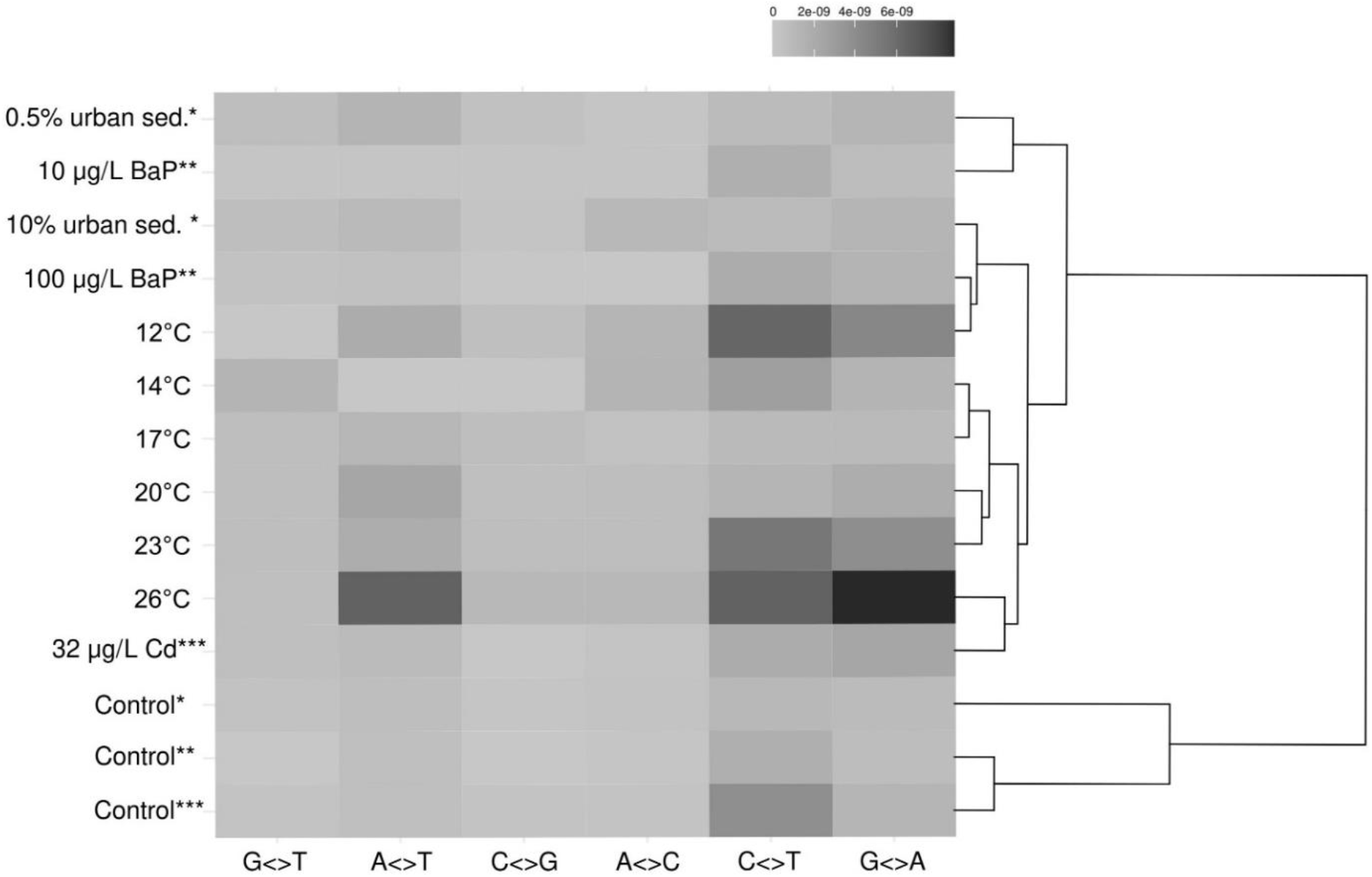
Heatmap and dendrogram of the mutational spectra induced on *C. riparius* by various environmental conditions.

## Discussion

While the substantial direct toxic effects of urban sediments containing TRWPs on *C. riparius* were recently evaluated (Rigano et al., 2024), the current study investigated a microevolutionary response of the same species to this stressor by directly assessing mutagenicity across multiple generations. This focus is underscored by the growing recognition of mutagenicity as a crucial endpoint in ecotoxicology, particularly in metazoans. Measuring genetic impacts is essential to fully understand the long-term ecological effects of contaminant exposure (Bulut et al., 2024; Doria and Pfenninger, 2021).

Both tested urban sediment concentrations (0.5% and 10%) are estimated to cause a 1.47- and 1.51-fold increase in mutation rates, respectively, compared to the control. This about 50% increase in mutagenicity highlights the potential of urban runoff sediments to cause lasting genetic changes, even at very low concentrations. Furthermore, given the pervasive presence of TRWPs in urban environments, such concentrations are likely common in surface waters (Baensch-Baltruschat et al., 2021; Mattsson et al., 2023; Zhang et al., 2014). The ecological relevance of *C. riparius* as a model organism in ecotoxicology suggests that our findings are significant not only for this species but may also be broadly applicable to other aquatic organisms. Consequently, sediment-dwelling freshwater biota in surface waters exposed to urban runoff likely experience persistent mutagenic effects, gradually altering the genetic composition of local populations.

Although mutations fuel evolution, most of them have deleterious effects (Crow, 2000). High mutation rates can, therefore, reduce individual fitness on average and pose a long-term risk to population stability. The accumulation of mutations can interfere with critical biological processes such as reproduction, development, and survival, ultimately threatening overall population viability (Charlesworth, 2009; Rigano et al., 2024a). Over time, accumulating mutations may paradoxically reduce genetic diversity (Lynch et al., 2016). This loss of diversity further compromises the population’s ability to adapt to environmental changes, increasing the likelihood of population decline or extinction (Lynch et al., 2016; Markert et al., 2010). However, we found no difference in mutation rate between the two concentrations, and thus no dose-response relationship.

The chemical characterization of the investigated sediment revealed a broad spectrum of chemical contamination related to traffic and urban impact. The measured tire rubber content was approximately 5 g/kg of dry sediment, with an estimated road runoff contribution of 4.5-8.5%, comparable to previous measurements from this sampling site (Rigano et al., 2024). The levels of salts and metals reflect the sediment accumulation period in the sedimentation basin, which spans late autumn and early winter, including freezing conditions. Road salts, the primary contributors to the overall contaminant load, not only induce osmotic stress but also accelerate the corrosion of vehicle parts, increasing the release of metals into road runoff. This effect is evidenced by the high concentrations of Fe and Al found in the matrix (Ebrahimi Gardeshi et al., 2024; Hintz and Relyea, 2019; Schuler and Relyea, 2018). Additionally, elevated concentrations of Pb, Cd, and Cu were detected, all linked to brake wear (Hwang et al., 2016; Rocha Vogel et al., 2024), with Cu being considered acutely toxic to aquatic wildlife (Brix et al., 2022) and phytotoxic in the mg/L range (Cruz et al., 2022). Furthermore, Cu toxicity in fish and invertebrates is closely linked to the salinity of the surrounding medium, suggesting that road salt loads may amplify the toxic effects of Cu and other metal contaminants in the environment (Grosell et al., 2007). However, little is known about the possible effects of co-exposure to Cd, Pb, and other metals (Balali-Mood et al., 2021; Liao et al., 2021; Wu et al., 2012; Yuan et al., 2014). Zinc in its oxidized form, zinc oxide (ZnO), is associated with oxidative stress responses in *C. riparius* and is commonly found in high concentrations as an additive in tire rubber (Gopalakrishnan Nair and Chung, 2015). The analysis of PAH revealed a dominant influence from high molecular weight (HMW) PAHs, suggesting a significant contribution from petrogenic sources (Tobiszewski and Namieśnik, 2012b). Furthermore, the presence of HMW PAHs, coupled with detected mineral oils, indicates a significant dioxin-like activity and mutagenic potential (Brinkmann et al., 2014; Grung et al., 2016; Shuliakevich et al., 2022). HMW PAHs may activate oxidative stress pathways, which could lead to mutagenic changes as a form of cellular adaptation to these toxic environmental conditions (Arambourou et al., 2019; Finckh et al., 2022). Concerning tire rubber additives, the detected organo-phosphate flame retardant Tris(1-chloro-2-propyl) phosphate (TCPP, 20 mg/kg dw) is known to cause DNA damage and act as a mutagen. However, little is known in general about the mutagenicity of tire rubber additives.

Previous studies have shown that ROS can cause oxidative damage to both chromosomal DNA and free nucleotides, leading to genetic alterations (Aitken and Krausz, 2001; Dröge, 2002; Sakai et al., 2006). Additionally, elevated ROS levels are often used as indicators of increased mutation rates (Aitken and Krausz, 2001). Our ROS analysis revealed, however, that at 0.5%, oxidative stress level increased significantly, while at 10%, ROS level was comparable to the control. Although it remains unclear why a lower concentration exerted higher oxidative stress, this pattern suggests that oxidative stress might primarily drive mutagenicity at lower concentrations, but not at higher ones. This interpretation is supported by the clustering of the respective mutation spectra in two distinct groups (Figure 4). While the spectrum induced by 0.5% urban sediment closely resembled that induced by 10 μg/L BaP, the spectrum induced by 10% urban sediment was like that induced by 100 μg/L BaP and low temperatures (12°C). This pattern suggested that different underlying mechanisms caused the mutations observed at the two concentrations of urban sediment and PAHs (polycyclic aromatic hydrocarbons). PAHs, which are abundant in complex mixtures like urban sediments (Rigano et al., 2024), are known to induce oxidative stress and DNA damage. Furthermore, BaP is considered a key marker of PAH exposure (Bukowska et al., 2022). However, a recent study discovered that pure BaP exposure actually reduced oxidative stress levels irrespective of the concentration, rather than inducing them (Bulut et al., 2024). However, this might be different in complex mixtures.

Thus, while oxidative stress cannot be excluded as the primary driver of mutagenicity at 0.5%, at higher concentrations like 10%, organisms may counteract ROS effects by activating detoxification pathways. These pathways employ cellular defenses to neutralize reactive oxygen species, such as the activation of superoxide dismutase (SOD) enzymes and catalase, converting them into less harmful by-products (Fukai and Ushio-Fukai, 2011). Despite the activation of these defense mechanisms, PAH-like compounds in urban sediments may still chemically modify the mutation spectrum, causing it to resemble that induced by 100 μg/L BaP. Alternatively, the observed consumption of urban sediment as a food source by the midges may explain the differences among concentrations. At 0.5%, the organic sediment content appeared to be mostly consumed by the larvae, whereas this was not the case at 10%. These organic molecules may capture most of the emerging ROS (Mir et al., 2024).

Exposure to low temperatures is necessarily associated with prolonged developmental times in insects due to reduced metabolic activity and slower cell cycle progression (Régnière et al., 2012). Likewise, the developmental time of *C. riparius* is significantly longer at 12°C (Oppold et al. 2016) and this might induce a higher mutation rate (Waldvogel and Pfenninger, 2021). The similarity of mutational spectra between low temperatures and 10% urban sediment suggested that the latter might have caused an increase in developmental time. However, previous studies have shown that exposure to BaP and urban sediments, especially at a concentration of 10%, does not prolong developmental time (Bulut et al., 2024; Rigano et al., 2024). This suggests that the similarities in mutation spectra observed at 10%, 100 μg/L BaP, and 12°C could result from either an unidentified common mechanism or coincidental effects.

Although heavy metals are a key component of urban sediments (Rigano et al., 2024, this study), the mutation spectrum did not resemble those of Cadmium (Doria and Pfenninger, 2021), which induces a mutation spectrum similar to those of very high temperatures (26°C). This is consistent with studies showing that heavy metals may act through mechanisms distinct from PAHs, including oxidative stress and interference with DNA repair processes (Chen et al., 2022).

The changes we observed in the mutational spectrum may have broader implications for genomic stability. The mutation spectrum plays a crucial role in shaping the AT content of genomes. Transitions and transversions from GC to AT, particularly in regions with high GC content, often lead to an overall increase in AT content. Normally, this shift is counterbalanced by GC-biased gene conversion (gBGC) during recombination, which preferentially preserves G and C bases (Capra et al., 2013; Mugal et al., 2015). However, the introduction of mutagenic agents, can disrupt this delicate equilibrium. In GC-rich genomic regions, changes in the mutation spectrum may result in a higher frequency of deleterious mutations and, therefore, reduced recombination efficiency (Capra et al., 2013). These changes could affect not only mutation rates but also the overall base composition of the genome, leading to long-term evolutionary consequences for populations exposed to these environmental stressors.

## Conclusions

The result of our study demonstrates that urban sediments containing TRWPs significantly increased the mutation rate of *C. riparius* by 50%, regardless of concentration. However, this was not the only mutagenic effect of urban runoff sediment. The observed changes of mutation spectra observed closely align with those induced by BaP, highlighting the putative role of PAH-like compounds in these genetic changes, which align with the considerable presence of PAHs, mineral oils, and tire rubber additives that have similar modes of action. Our results indicated that urban sediments not only impacted individual fitness but also reshaped genetic diversity and evolutionary dynamics. This highlights the need for multigenerational studies to better understand and manage the long-term effects of complex mixtures of pollutants and urban impacts on natural populations and entire ecosystems.

## Supporting information

Supplementary Infos

## Acknowledgments

The study received support from the LOEWE-Centre Translational Biodiversity Genomics (TBG) (LOEWE/1/10/519/03/03.001(0014)/52) funded by Hessen State Ministry of Higher Education, Research and the Arts (HMWK). The RoadTox project was funded by the Ministry of the Environment, Nature and Transport of the State of North Rhine-Westphalia. The LC-HRMS instrument used for target screening analysis at the Helmholtz Centre for Environmental Research is part of the major infrastructure initiative CITEPro (Chemicals in the Environment Profiler) funded by the Helmholtz Association.

